# Adaptation to Solvent Environment in Toll-like Receptor 5: A Comparative Evolutionary Analysis of Membrane-bound and Soluble Forms in *Epinephelus coioides*

**DOI:** 10.1101/2025.02.28.640895

**Authors:** Alper Karagöl, Taner Karagöl

**Author notes:** To whom the correspondence should be addressed. Email: Alper Karagöl, Taner Karagöl. These authors contribute equally.

## Abstract

Toll-like receptor 5 (TLR5) is a conserved member of the innate immune system and known for its ability to detect bacterial flagellin. Interestingly, some teleost fish, including the orange-spotted grouper (*Epinephelus coioides*), possess both membrane-bound (mTLR5) and soluble (sTLR5) forms of TLR5, which is not observed in mammals. This study investigates the evolutionary patterns towards adapting a solvent environment in TLR5 by comparing the membrane-bound (mTLR5) and soluble (sTLR5) forms from the *Epinephelus coioides*. We analyzed sequence characteristics, amino acid composition, and physicochemical properties of both proteins to understand the basis for their differing solubility and polarity. Two proteins share a substantial part of structural and evolutionary characteristics. Superposition of Alphafold3 predicted structures quantified the structural similarities with a RMSD value of 1.361Å. Contrary to the conserved tertiary structures, our results reveal distinct amino acid preferences between sTLR5 and mTLR5, suggesting that specific mutations may have driven the evolution of the soluble form. Asymmetric evolutionary dynamics and kernel causality were thereby investigated using generalized correlation coefficients (gmc). The dependence analysis revealed that TLR5 evolution shows multidirectional dynamics between soluble and membrane-bound forms, as 2 amino acids (M and V) shift membrane-to-solvent, 2 shift solvent-to-membrane (H and Y). The T<=>V and D>K substitutions required two base changes and therefore introduced a mutational bias. Regression analysis of the homologous sequences indicated T<=>V changes may have supported by alanine intermediates. These findings provide insights into the functional diversification of TLR5 and broader implications on the diversification of immune system proteins.

**Graphical Abstract.**
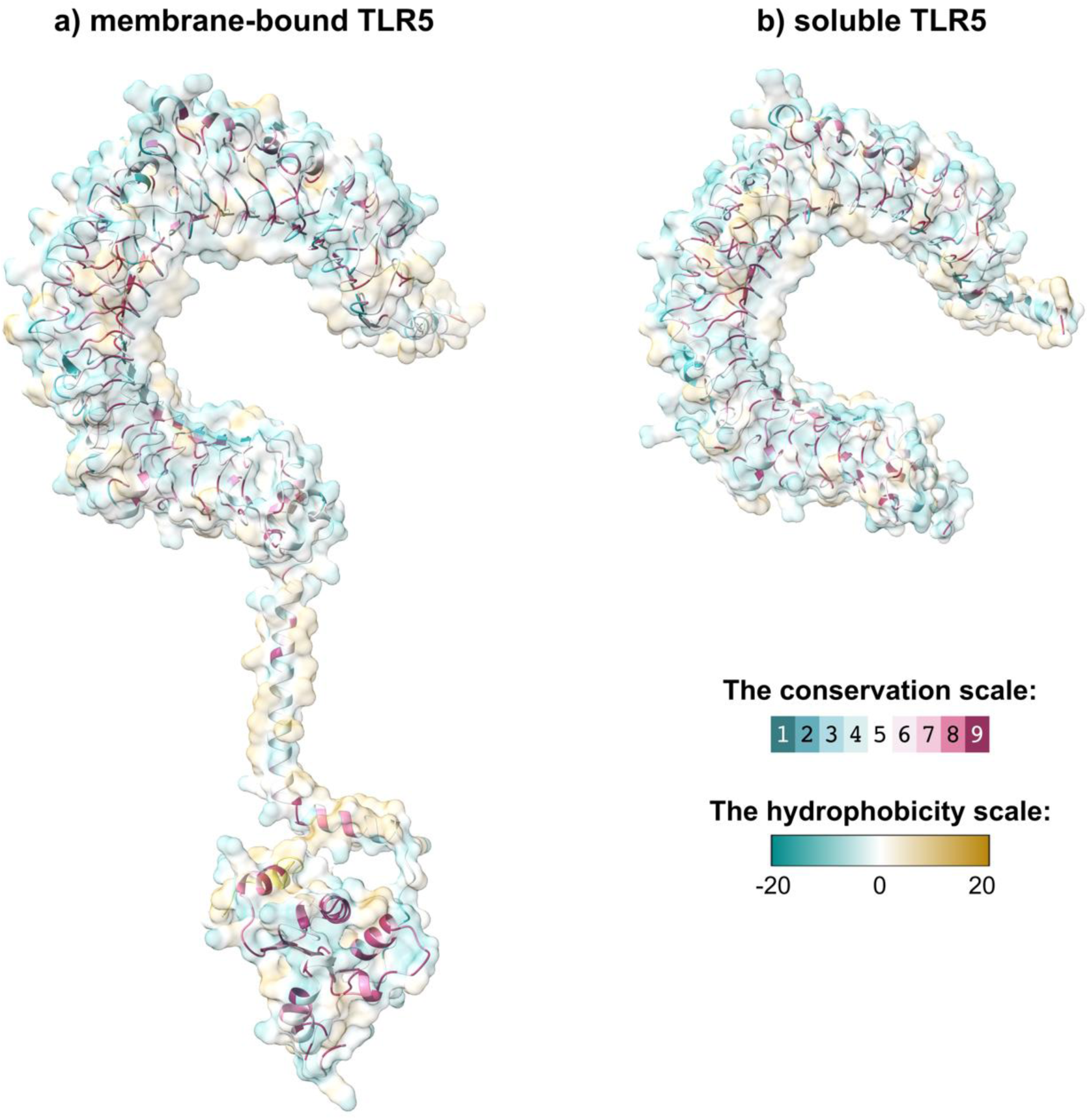
Surface profiles and evolutionary conservation of membrane-bound and soluble TLR5. Molecular surface representations were coloured by hydrophobicity (cyan: hydrophilic, gold: hydrophobic). The underlying colour map represents the conservation grades of two proteins. Evolutionary conservation grades of each amino acid residue predicted by Consurf server; visualized by the color-coding scheme of nine colours, ranging from turquoise (variable) through white (average) through burgundy (conserved) represents conservation grades 1 to 9, in order of increasing conservation (1= Variable, 5= Average, 9= Conserved).

## Introduction

<u>T</u>oll-like receptors (TLRs) play a fundamental role in the innate immune system, recognizing pathogen-associated molecular patterns and initiating immune responses (1). Among these, TLR5 is known for its ability to detect bacterial flagellin (1, 2, 3). Interestingly, some teleost fish, including the orange-spotted grouper (*Epinephelus coioides*), possess both <u>m</u>embrane-bound (mTLR5) and <u>s</u>oluble (sTLR5) forms of TLR5, which is not observed in mammals (4) The membrane form of TLR5 (mTLR5) shares a similar structure with mammalian TLR5 (4, 5). Meanwhile, sTLR5 is primarily composed of the mTLR5 ectodomain that contains leucine-rich repeats (LRR) (4). The stability of the horseshoe-like structure of the TLR5 molecule depends on the rigidity of LRR folding and spatial configuration (6).

Soluble proteins can diffuse more freely in cellular environments, potentially allowing for more diverse functional outcomes (7). In the case of sTLR5, this solubility may result in advantages for pathogen detection and immune response initiation. The soluble protein may function as a negative modulator that contribute to TLR signaling pathways by antagonizing the proinflammatory activity of mTLR, as such roles were previously identified for TLR2 (8, 9). Both sTLR5 and mTLR5 were widely expressed in the liver and immune-related organs and tissues, but they exhibited distinct expression profiles (4). Concurrently, the exact roles of sTLR5 in the regulation of flagellar-mediated immune responses remain inconclusive. Recent studies have explored the structural and functional differences between mTLR5 and sTLR5 in various fish species (4, 5). Accordingly, the functional domains of TLRs showed a significant conservation, with variations primarily observed in the N-terminal region of the LRR domain (5). On the other hand, the adaptative changes specifically for the solvent environment remained largely unstudied.

The evolution of protein solubility is a fundamental aspect of protein function and adaptation (7). The constraints and plasticity in the adaptation to soluble environments is fundamental and crucial, and this specific case of TLR5 evolution provides a useful framework as the presence of both membrane-bound and soluble forms of TLR5 in teleost fish raises questions about the evolutionary pressures that led to this divergence. On the other hand, the transition is unlikely to be unidirectional (membrane to soluble), instead possibly multidirectional and complex. In this case, evolutionary analysis and variational profiling could have a strong potential to explain the functional implications of these chemically distinct forms. Additionally, the absence of transmembrane and intracellular domains in sTLR5 necessitates a reorganization of its tertiary structure to maintain functional integrity, potentially involving the formation of new intramolecular interactions to compensate for the loss of membrane-associated stabilizing factors. Understanding the evolutionary shifts from solvent to membrane environments and vice versa in TLR5 provides a framework for designing synthetic receptors that can adapt to different cellular compartments or environmental conditions. Understanding natural variations could inform the design of proteins with enhanced properties, including solubility (10). Interestingly, our previous analyses suggest despite its high mutational barrier, T<=> V substitutions were less harmful than other polar to nonpolar changes (10, 11). However, these substitutions were not seen in human single nucleotide variation databases, unlike other polarity altering substitutions of I<=>T, F<=>Y, and L<=>Q (10, 11). This study further identifies 4 natural T<=>V changes in aligned residues, closing a significant gap in our understanding of adaptation to solvent environments. We suggest utilizing variations with higher mutational barrier, especially acidic to basic (D>K) and polar to non-polar (T>V) changes, is potentially useful in analysing selective pressures. Our study further embraces the complexity of molecular evolution by including a modular approach, where some protein regions largely coupled or more sensitive to mutational changes, while others evolve more neutrally.

In this study, we aim to investigate the molecular basis for the evolution of mTLR5 and sTLR5 in *E. coioides*. We hypothesize that specific changes in amino acid composition and physicochemical properties have contributed to the diversification of the soluble form from its membrane-bound counterpart. By comparing the sequences and characteristics of mTLR5 and sTLR5, we seek to identify key features that may have driven this evolutionary transition and to understand the potential functional implications of these changes.

## Results and Discussions

### Psychochemical properties of mTLR5 and sTLR5

The mTLR5 sequence consists of 821 amino acids, while sTLR5 is shorter with 644 amino acids (4, 5). This is because the soluble protein lacks the transmembrane domain seen in the membranous form (4). The structure of TLRs is characterized by a N-terminal ectodomain made of leucine-rich repeats, a characteristic solenoid structure giving to the ectodomain a horseshoe-shape, a single transmembrane domain, and a C-terminal intracellular signalling domain (4, 5, 6). Although they are important for function (12), this study focuses on structural aspects of two proteins, so it specially analyses the difference between matched positions between sTLR5 and extracellular aligned domain of mTLR5.

Sequence alignment revealed a 56.4% identity in 626 aligned residues (Figure 1). The most notable difference is the absence of the transmembrane and intracellular domains in sTLR5, consistent with its soluble nature. The transmembrane helix in mTLR5 between residues 646^th^ and 668^th^ is also detected by DeepTMHMM, a deep learning model for transmembrane topology prediction (13). The increased number of charged residues in sTLR5 likely enhances its hydrophilicity and solubility. mTLR5 has 53 negatively charged, 51 positively charged residues in extracellular aligned part; meanwhile sTLR5 has 58 negatively charged, 54 positively charged residues. The negative GRAVY score of sTLR5 indicates a higher hydrophilicity compared to mTLR5, consistent with its soluble nature. GRAVY score for mTLR5 was 0.016, and for sTLR5 it was −0.035. The slightly higher aliphatic index of sTLR5 suggests increased thermostability, which may contribute to its solubility. Both proteins are classified as stable, with sTLR5 showing a slightly higher instability index, mTLR5: 37.29 (stable); sTLR5: 39.75 (stable).

**Figure 1.**
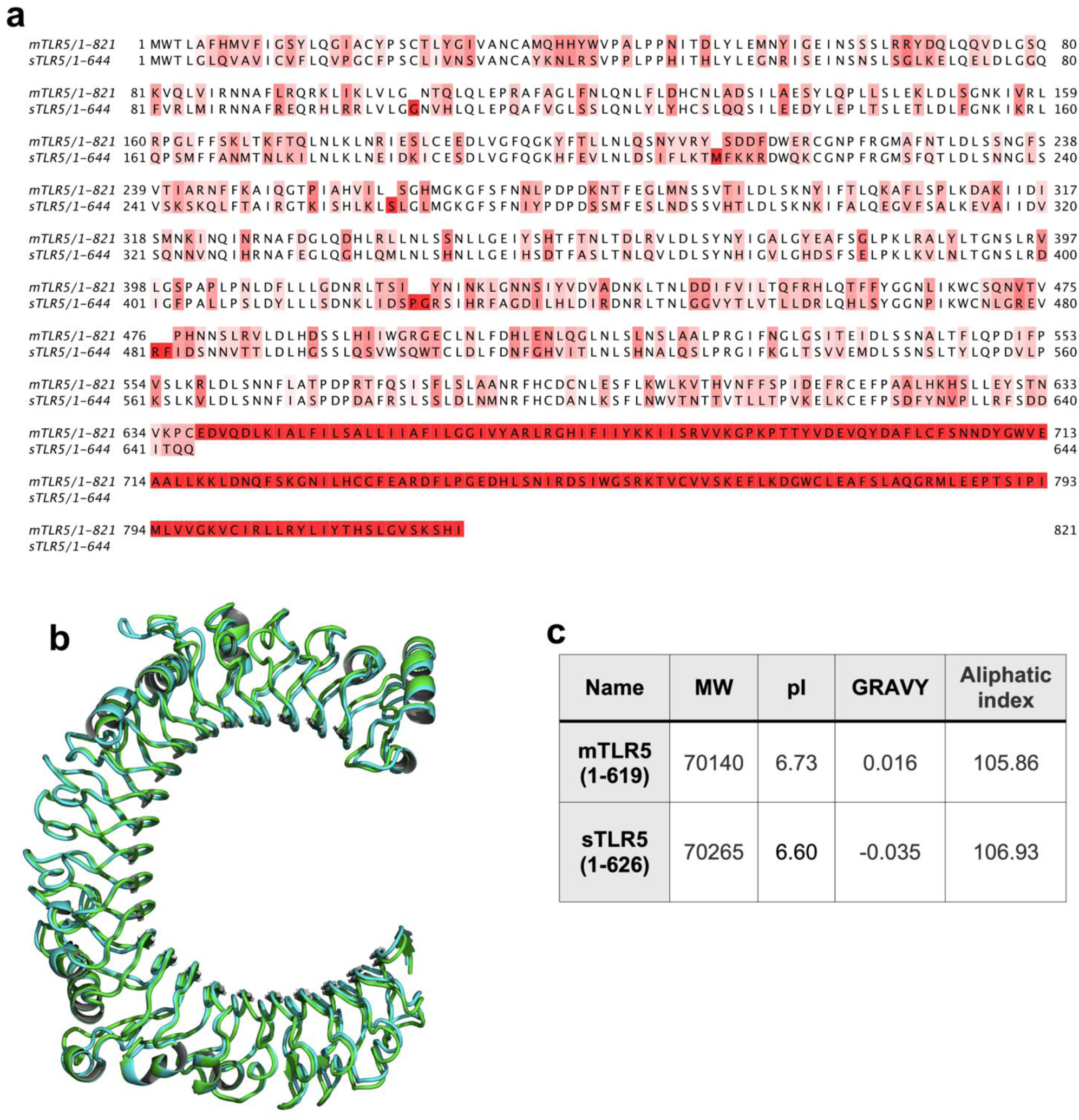
Sequence alignment (a) and superimposed structures (b) of mTLR5 and sTLR5 that are predicted by AlphaFold3. Structural data for mTLR5 (green) and sTLR5 (cyan). The root <u>m</u>ean <u>s</u>quare <u>d</u>eviation (RMSD) values of 1.361Å. N-termini loops are removed to facilitate direct comparisons.

### Amino Acid Composition Analysis

Comparative analysis of amino acid composition revealed several key differences between mTLR5 and sTLR5. Firstly, aligned portion of the sTLR5 showed higher percentages of valine (V), serine (S), histidine (H), and glutamic acid (E) compared to mTLR5. Charged residues on the protein surface can create electrostatic repulsion between protein molecules (14, 15). This repulsion helps prevent unwanted protein-protein interactions and aggregation, which is essential for a soluble protein circulating in extracellular fluids (15).

mTLR5 had higher percentages of alanine (A), tyrosine (Y), and glutamine (Q) compared to sTLR5 alignment. This may indicate an adaptation to membrane environments. Alanine (A) is small and can pack well in membrane proteins, Q and Y are polar. Their increase in mTLR5 might facilitate interactions with other cell surface molecules or contribute to the flagellin binding site. Both forms had high percentages of leucine (L), with sTLR5 slightly higher (17.6%) than mTLR5 (17.0%). Leucine is already highly represented in both forms, this suggests leucine-rich repeats is preserved for the protein’s structure or function, making dramatic changes of leucine count less likely (4). These differences suggest a shift towards amino acids that favour solubility in sTLR5, particularly the increase in polar (S) and charged (E, H) residues.

### Residue-wise substitutions

While L percentages remain almost unchanged, the most frequent substitution was F>L (phenylalanine to leucine), occurring 10 times and resulting in a change from a nonpolar aromatic to a nonpolar aliphatic residue, with an increase of 1 on the Kyte-Doolittle hydrophobicity scale (Table 1). The change to L could be related to Leucine rich domain. Four substitutions (I>V, N>S, R>K, and A>S) were each observed 7 or 6 times. Notable among these was the A>S change, which resulted in a large decrease in hydrophobicity (−2.6) among all observed substitutions. I>V maintained the nonpolar aliphatic nature but slightly decreased hydrophobicity by 0.3. N>S involved a change from a polar amide to a polar hydroxyl group, increasing hydrophobicity by 2.7.

**Table 1.**
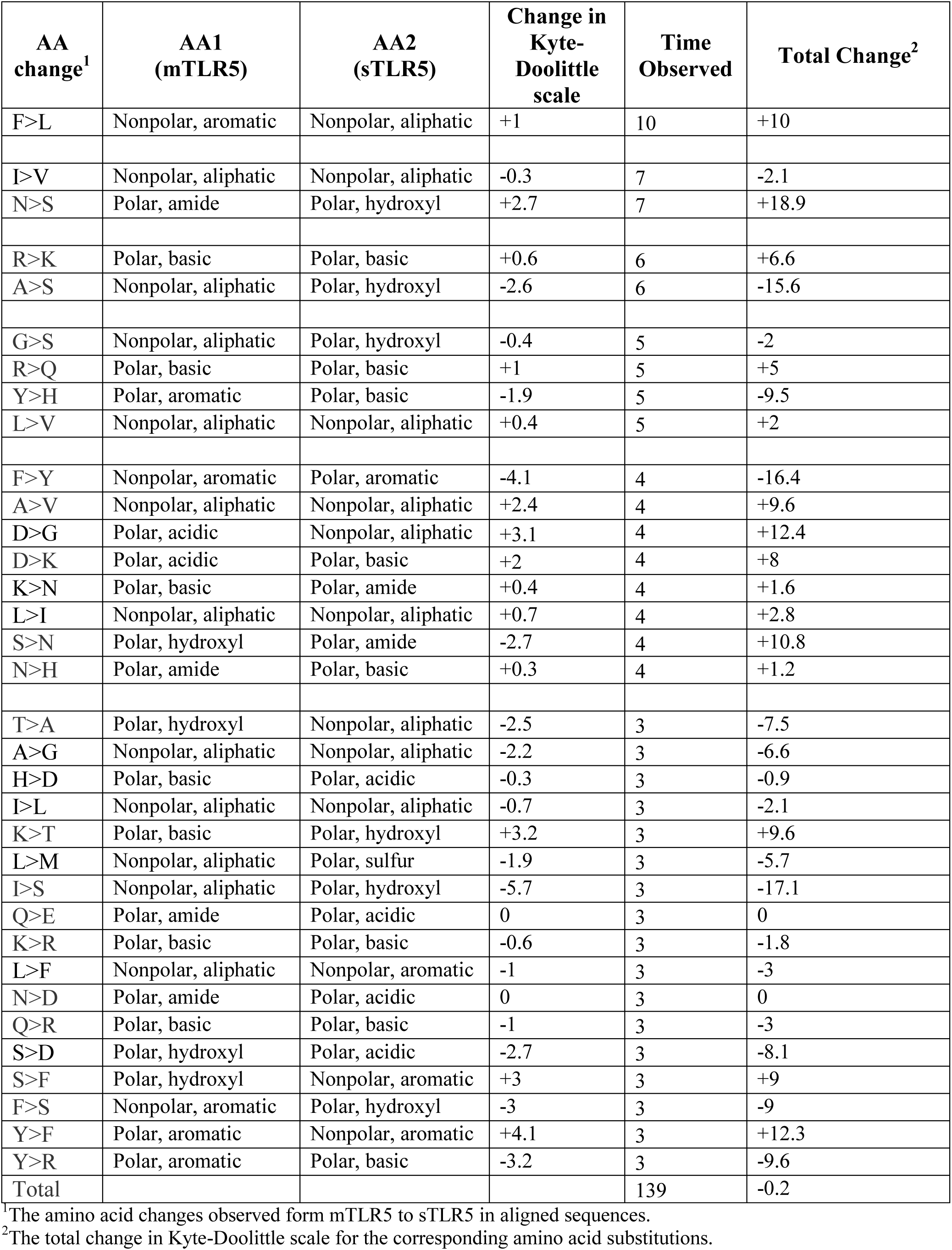
Substitutions observed more than 2 times.

Four substitutions (G>S, R>Q, Y>H, and L>V) occurred 5 times each. The G>S change involved a shift from a nonpolar aliphatic to a polar hydroxyl group, decreasing hydrophobicity by 0.4. Eight substitutions (F>Y, A>V, D>G, D>K, K>N, L>I, S>N, and N>H) were observed 4 times each. The F>Y substitution caused the largest decrease in hydrophobicity (−4.1) among these changes, while D>G led to a substantial increase in hydrophobicity (+2.4).

Overall, the amino acid changes involved various shifts between nonpolar, polar, aromatic, aliphatic, acidic, and basic residues. The changes in hydrophobicity ranged from −4.1 to +3.2 on the Kyte-Doolittle scale, indicating significant alterations in the protein’s hydrophobic properties. The remaining 17 substitutions occurred 3 times each. Total hydrophobicity changes from these substitutions were −0.2, where sums of −16.4 and −17.1, and −15.6 comes from F>Y, I>S and A>S substitutions, respectively. Interestingly, the F>Y was included in the QTY-code (10, 16) On the other hand, the effect of these substations on polarity remained subtle (−0.2), indicating that rather than a quantity-oriented profile, rare substitutions resulted the overall increase in polarity in the sTLR5.

Several changes involve amino acids commonly subject to post-translational modifications, such as serine and lysine (17, 18). N>S and A>S (increase in serine); Y>H and Y>F (decrease in tyrosine). There are many substitutions occurred that involves serine residues. Interestingly, serine is unique among amino acids in being encoded by two non-overlapping codon sets (TCN and AGY), these distinct codon sets drive different substitution patterns at sites under purifying and diversifying selection pressures (19). K>N and K>T (decrease in lysine) were also shown, lysine can be subject to various modifications like acetylation, methylation, and ubiquitination (18). These changes could affect the regulation of the protein’s activity, stability, or interactions with other molecules.

This bidirectional nature of changes suggests several possibilities (Table 2). The reversibility may indicate that these positions are not under strong selective pressure, allowing for some variability without severely impacting function. Some of these changes may be functionally neutral, persisting due to random genetic drift rather than selective pressure (20). Many changes (like F<=>L, I<=>V) involve amino acids with similar properties, suggesting a conservation of overall function while allowing for some variability. Conversely, these substitutions were not strongly reversible (the directions were predominantly F>L, I>V). Direction preference suggests selection pressures beyond mutation probability, indicating the importance of considering directional pathways in a correlation analysis. Some of the changes are considered irreversible because they were only observed in one direction (Table 2). For example, Y>H was observed 5 times, but there were no instances of H>Y. Some of these changes represent significant shifts in amino acid properties: From aromatic to basic (Y>H) and from acidic to basic (D>K). D>K also requires at least 2 substantial base changes in same codon: Aspartic acid (D) GAT, GAC; and Lysine (K) AAA, AAG. This may result in making the reversible substitution less likely.

**Table 2.** Reversible substitutions.

| AA change | Codon change<sup>1</sup> | mTLR5>sTLR5<sup>2</sup> | sTLR5>mTLR5<sup>3</sup> | Total<sup>4</sup> | Net Change<sup>5</sup> |
| --- | --- | --- | --- | --- | --- |
| F<=>L | U<=>C (1st base) | 10 | 2 | 12 | 8 |
| I<=>V | A<=>G (1st base) | 7 | 1 | 8 | 6 |
| A<=>S | G<=>U (1st base) | 6 | 2 | 8 | 4 |
| G<=>S | G<=>A (1st base) | 5 | 1 | 6 | 4 |
| N<=>S | A<=>G (2nd base) | 7 | 4 | 11 | 3 |
| R<=>K | G<=>A (2nd base) | 6 | 3 | 9 | 3 |
| N<=>H | A<=>C (1st base) | 4 | 2 | 6 | 2 |
| L<=>V | U<=>G (1st base) | 5 | 4 | 9 | 1 |
| F<=>Y | U<=>A (2nd base) | 4 | 3 | 7 | 1 |
| L<=>I | C<=>A (1st base) | 4 | 3 | 7 | 1 |
| F<=>S | U<=>C (2nd base) | 3 | 3 | 6 | 0 |
| Y>H | U<=>C (1st base) | 5 | 0 | 5 | 5 |
| A<=>V | C<=>U (2nd base) | 4 | 1 | 5 | 3 |
| K<=>N | A<=>G (3rd base) | 4 | 1 | 5 | 3 |
| D>K | G<=>A (1st base) &<br>U/C <=>A/G (3rd<br>base) | 4 | 0 | 4 | 4 |
| D>G | A<=>G (2nd base) | 4 | 0 | 4 | 4 |
| A<=>G | C<=>G (2nd base) | 3 | 1 | 4 | 2 |
| I<=>S | U<=>G (2nd base) | 3 | 1 | 4 | 2 |
| K<=>T | A<=>C (2nd base) | 3 | 1 | 4 | 2 |
| N<=>D | A<=>G (1st base) | 3 | 1 | 4 | 2 |
| Q<=>E | C<=>G (1st base) | 3 | 1 | 4 | 2 |
| D<=>E | U<=>G (3rd base) | 2 | 2 | 4 | 0 |
| T<=>V | A<=>G (1st base) &<br>C<=>G (2nd base) | 2 | 2 | 4 | 0 |

<sup>1</sup>The nucleotide base change associated with the amino acid substitution.
<sup>2</sup>The number of occurrences where the change observed form mTLR5 to sTLR5 in aligned sequences.
<sup>3</sup>The number of occurrences where the change observed form sTLR5 to mTLR5 in aligned sequences.
<sup>4</sup>The total number of changes for the corresponding amino acid substitution.
<sup>5</sup>The absolute difference between mTLR5 > sTLR5 and sTLR5 > mTLR5.

Another less likely change is T<=>V, threonine (T) codons: ACU, ACC, ACA, ACG; valine (V) codons: GUU, GUC, GUA, GUG. Thereby, a transition between these amino acids requires at least 2 base changes. Interestingly, our previous analyses suggest despite its high mutational barrier, these substitutions were less harming than other polar to nonpolar changes (10, 11). This study further identifies 4 T<=>V substitutions in the dataset, that were not possible in human single nucleotide variation databases (10, 11). The direction of these amino acid changes is another significant observation, no dominant direction was found, as 2 T>V and 2 V>T substitutions were observed. Although the sample size was small, acidic to basic (D>K) and polar to non-polar (T>V) changes potentially useful in analysing selection pressures.

### Qualitative vs. Quantitative Mutational Changes and Extensive Forms

While the proposed diversification of sTLR5 from mTLR5 involves some quantitative changes in amino acid composition, our analysis reveals that the mutations go beyond simple quantity-based alterations. Despite the substantial number of substitutions, the overall structures remained similar as RMSD value of superposed structures was 1.36 Å (Figure 1). The mutations were not uniformly distributed throughout the protein. Instead of merely increasing or decreasing the quantity of certain amino acids, we observed strategic substitutions that enhance solubility while maintaining structural integrity. For example, the increase in valine (V) in sTLR5 is accompanied by a decrease in isoleucine (I), both hydrophobic residues. This substitution likely maintains necessary hydrophobic interactions while potentially increasing flexibility. The higher proportion of serine (S) in sTLR5 is balanced by a decrease in threonine (T), both polar and hydroxyl residues. This change may fine-tune the hydrophilicity without drastically altering the overall structure. Interestingly, the valine (V), isoleucine (I) and threonine (T) residues were also important in QTY-code protein design approach (16). It was previously demonstrated that evolutionary T<=>V changes could result from intermediate alanine residues (11). For instance, T>A (3 times) and A>V (4 times) changes occur in separate residues, increasing the number of valine and decreasing threonine (Table 1). For our dataset, 2 T>V and 2 V>T substitutions were observed (Table 2). Apparently, the T>V changes occur in number but a more indirect way, a direct change requires two base changes in same codon and unlikely in nature. In the case of these indirect substitutions, the evolution of polarity changes of mTLR5 to sTLR5 can be modelled as a sequence of evolutionary adaptations. Extensive form games can be used to represent the sequential nature of these evolutionary changes, where each adaptation (e.g., transition from membrane-bound to soluble form) affects subsequent evolutionary opportunities and constraints (21). In this case, the intermediate alanine residue in a T>V variation could represent a specific decision node in the evolutionary pathway.

### Co-evolution of residues

The similarity in the overall shape and the distribution of conserved regions between the two molecules suggest roles of certain structural elements in TLR5 across different variants, while also displaying areas where evolutionary flexibility is more permitted (Figure 2). The <u>e</u>volutionary <u>c</u>oupling (EC) analysis suggests a differential evolutionary pressure on different parts of the protein. The frequently co-evolved residues predominantly appear in the concave face of the horseshoe (Figure 2). The conserved regions likely to indicate critical functions such as flagellin recognition, while the more variable regions may adapt to diverse environmental pressures or facilitate species-specific interactions. The observed evolutionary coupling may have significant implications for TLR5 function and adaptation. It suggests a modular approach to protein evolution, where core recognition elements remain largely coupled or sensitive to mutational changes across protein, while peripheral regions evolve more neutrally. This strategy could enable TLR5 to maintain its essential role in innate immunity while adapting to environment-specific requirements.

**Figure 2.**
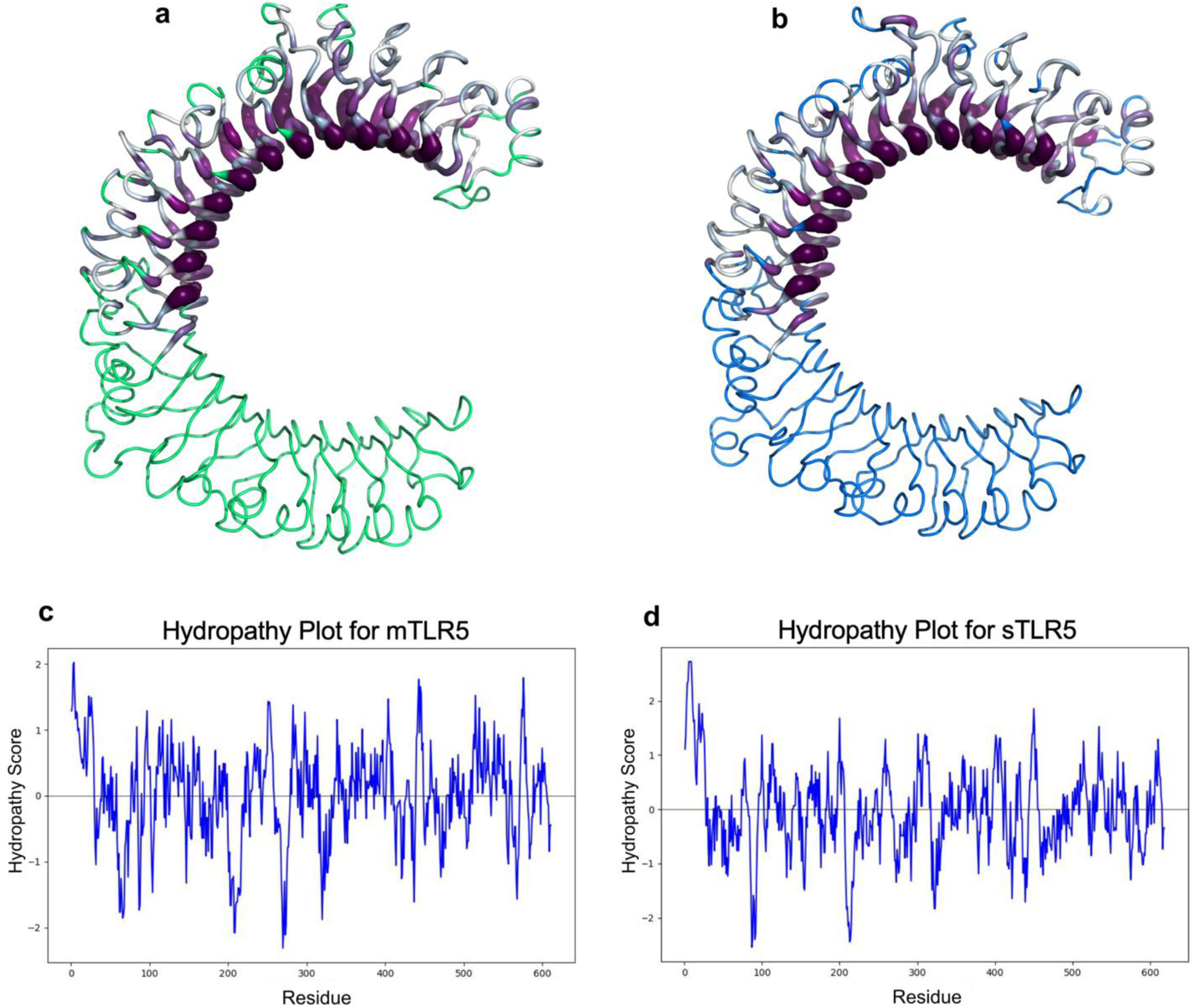
Evolutionary coupling dynamics of mTLR5 (a) and sTLR5(b) and Kyte-Doolittle hydropathy plots (c and d). (a) mTLR5 structure with evolutionarily coupled nodes depicted in purple, contrasted with regions of lower coupling frequency in green. (b) sTLR5 structure emphasizing the frequently coupled nodes (purple) and lower coupled (cyan) domains. Both structures exhibit the characteristic horseshoe-shaped leucine-rich <u>r</u>epeat (LRR) domain topology typical of Toll-like receptors. (c and d) Residue-wise Kyte-Doolittle hydrophobicity plots for mTLR5 and sTLR5, respectively.

Both proteins demonstrate fluctuating residue-wise hydropathy scores across approximately 600 residues. However, in C-terminal region (residues 500-600), mTLR5 shows higher variability with several pronounced peaks and troughs. Meanwhile, sTLR5 displays a more moderate pattern with less extreme fluctuations. Interestingly, both proteins show significant hydrophilic regions (negative Kyte-Doolittle scores), but mTLR5 has deeper troughs. sTLR5, as a soluble form, could show adaptations for functioning independently in extracellular fluids. This is evident in the smoother transitions between hydrophobic and hydrophilic regions, which indicates a more uniformly folded structure and less dramatic changes in its surface properties (Figure 2). These distinct hydropathy profiles suggest while there are notable similarities, mTLR5 and sTLR5 may have evolved different structural features to support divergent cellular localizations.

The analysis of specific amino acid frequencies in evolutionary data produces insights into the evolutionary dynamics of hydrophobic and hydrophilic residues in sTLR5 (Table 3). Valine (V) and threonine (T) frequencies were not statistically significant. However, when V and T frequencies were regressed on alanine (A), the directionality showed a strong kernel causal influence of valine leading to threonine (V>T) with significant correlation (0.19, p < 0.001). This suggests that valine’s hydrophobic nature plays a critical role, with selective pressure toward threonine under certain evolutionary contexts, likely where polar properties are favored.

**Table 3.** Evolutionary profiles of the sampled substitutions from sTLR5.

| AA change | Partial Pearson (AA1, AA2)<sup>1</sup> | Partial Correlation (AA1\|AA2)<sup>2</sup> | Partial Correlation (AA2\|AA1)<sup>3</sup> | Kernel Cause<sup>4</sup> | Causal Direction<sup>5</sup> |
| --- | --- | --- | --- | --- | --- |
| V<=>T | 0.061<br>(p>0.05) | 0.017<br>(p>0.05) | 0.002<br>(p>0.05) | - | - |
| V<=>T<br>(regressed on A)<sup>6</sup> | 0.069<br>(p>0.05) | 0.034<br>(p>0.05) | 0.19<br>(p<0.001) | V | V>T |
| F<=>Y | 0.083<br>(p>0.05) | 0.125<br>(p<0.001) | 0.26<br>(p<0.001) | F | F>Y |
| I<=>V | 0.235<br>(p<0.001) | 0.164<br>(p<0.001) | 0.299<br>(p<0.001) | I | I>V |
| N<=>S | -0.045<br>(p>0.05) | -0.126<br>(p=0.002) | -0.001<br>(p>0.05) | S | S>N (-) |
| R<=>K | 0.223<br>(p<0.001) | 0.157<br>(p<0.001) | 0.186<br>(p<0.001) | R | R>K |
| R<=>Q | 0.187<br>(p<0.001) | 0.343<br>(p<0.001) | 0.113<br>(p=0.005) | Q | Q>K |
| G<=>S | -0.007<br>(p>0.05) | 0.149<br>(p=0.002) | 0.247<br>(p<0.001) | G | G>S |
| D<=>G | -0.008<br>(p>0.05) | -0.181<br>(p<0.001) | -0.036<br>(p>0.05) | G | G>D (-) |
| S<=>N | -0.045<br>(p>0.05) | -0.008<br>(p>0.05) | -0.127<br>(p=0.002) | S | S>N (-) |

<sup>1</sup>Partial Pearson correlation, controlling for evolutionary conservation of the residue (Consurf Grade).
<sup>2</sup>Partial generalized correlation, where AA2 (sTLR5) is the kernel cause, controlling for evolutionary conservation of the residue (Consurf Grade).
<sup>3</sup>Partial generalized correlation, where AA1 (mTLR5) is the kernel cause, controlling for evolutionary conservation of the residue (Consurf Grade).
<sup>4</sup>Kernel causality refers to a class of methods that employ kernel functions to model causal relationships between variables in a non-parametric framework. If absolute value of Partial generalized Correlation (AA1|AA2) is bigger than (AA2|AA1), AA2 is said to be the kernel cause, or vice versa. Threshold = 0.02.
<sup>5</sup>Causal direction according to kernel cause.

The substitution between phenylalanine (F) and tyrosine (Y), both aromatic residues, shows a strong evolutionary trend with F>Y (0.125 and 0.26, p < 0.001). The substitutions involving charged residues, such as arginine (R) to lysine (K), show a significant and stronger causal direction (R>K, 0.157 and 0.186, p < 0.001). The close similarity in the positive charge of these residues suggests that evolutionary changes maintain the overall charge while potentially fine-tuning protein interactions. The transition from R to K could imply a preference for lysine’s smaller size in certain regions of sTLR5, allowing for more flexible interactions or structural adjustments.

In another case, the substitutions demonstrate a significant directional change from glutamine to glycine (Q>K, 0.343 and 0.113, p < 0.001). This indicates an evolutionary shift where the neutral-polar residue (Q) may occasionally be replaced by a charged residue (K). Interestingly, the substitution between aspartate (D) and glycine (G) shows a negative correlation (G>D, −0.181, p < 0.001), indicating that glycine might be favoured over aspartate in certain regions possibly due to its flexibility and ability for a greater conformational freedom. These adaptation to solvent environment may allow for more diverse functional roles of sTLR5 in the immune system of teleost fish, potentially enhancing their ability to respond to bacterial flagellin in different cellular compartments.

### Kernel Causality and Asymmetric Evolutionary Dynamics of TLR5

The observed differences in amino acid composition and physicochemical properties between mTLR5 and sTLR5 suggest an evolutionary trend towards increased solubility in sTLR5. On the other hand, evolutionary changes could be multidirectional. The selection of sTLR5 and mTLR5 may show themselves favouring changes both for membrane bound and soluble forms. The conservation score of 0.950 suggests that while there are some differences between sTLR5 and mTLR5, they maintain a high degree of similarity, likely due to shared functional requirements (Supplementary Figure S1, Figure S2). A, C, D, E, F, G, I, K, L, N, P, Q, S, and W show high conservation between sTLR5 and mTLR5 (Pearson correlation > 0.98), suggesting crucial roles in maintaining overall structure and function.

The dynamic between the TLR5 soluble form (sTLR5) and the membrane-bound extracellular domain (mTLR5) is asymmetric, rather than a focused adaptation to solvent environment. H and Y show a tendency to move from solvent to membrane in TLR5, suggesting potential adaptations for surface interactions in the membrane-bound form (Table 4). Interestingly, Y was shown stronger unidirectional potential compared to H. This may reflect a compensatory adaptation where mTLR5 integrates characteristics necessary for membrane stability.

**Table 4.**
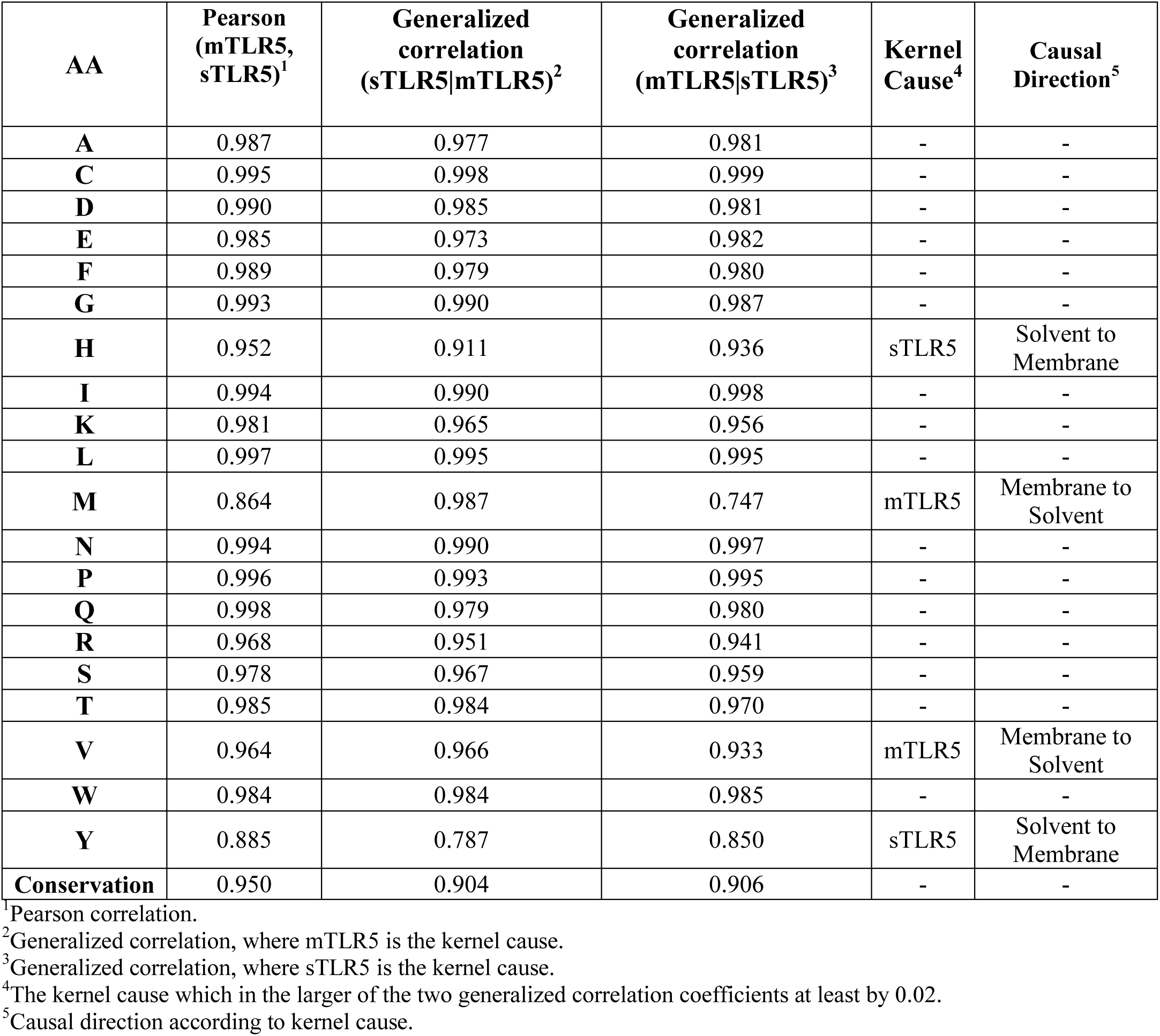
Residue-wise evolutionary dynamics of sTLR5 and mTLR5.

M, and V show a tendency to move from membrane to solvent, possibly indicating adaptive changes for ligand binding in the soluble form. Notably, M (0.864) shows the lowest Pearson correlation, with a strong tendency to move from membrane to solvent in mTLR5. The low generalized correlation (mTLR5|sTLR5) value for M (0.747) suggests that the positions of M residues in mTLR5 are not exact predictors of M positions in sTLR5, which would be expected if alternative splicing altered the N-terminal region. It is important to note that other mechanisms, such as gene duplication followed by divergence, could potentially produce similar patterns. This suggests that while sTLR5 maintains characteristics of mTLR5, the reverse is not as strongly true, indicating some unidirectional adaptation for the solvent environment.

### Codon Usage Bias

The evolution of sTLR5 involves a redistribution of charged residues rather than a simple increase. While the total number of charged residues increases slightly, we see a more significant shift in the ratio of negatively to positively charged residues, potentially optimizing electrostatic interactions for solubility. Analysis of the coding sequences might reveal changes in codon usage between mTLR5 and sTLR5. Such changes could affect translation efficiency and protein folding, contributing to solubility without altering the amino acid sequence (22). While both TLR5 variants show some level of codon usage optimization for expression in *Danio rerio* (zebrafish), there is potential for further optimization, particularly in terms of tRNA adaptation and codon pair usage. Both genes showed moderate codon usage bias, with <u>e</u>ffective <u>n</u>umber of <u>c</u>odons (ENC) values of 54.57 for mTLR5 and 53.69 for sTLR5 (range: 20-61). Both genes exhibited similar <u>e</u>ffective <u>n</u>umber of <u>c</u>odon <u>p</u>airs (ENcp) values: 35.19 for mTLR5 and 35.87 for sTLR5. The <u>c</u>odon <u>a</u>daptation index (CAI) was slightly higher for sTLR5 (0.75) compared to mTLR5 (0.71), indicating potentially better adaptation to the host’s preferred codons.

### Future scopes and the potential applications

The removal of the transmembrane and intracellular domains in sTLR5 is accompanied by specific changes in the remaining sequence, also showed themselves in the co-evolution data of aligned residues. Our study reveals bidirectional changes and their roles in molecular diversification of TLR5. This approach could be used with advanced statistical tools such as kernel regression and stochastic dominance analysis to more accurately represent and analyze specific variations. These transitions, particularly in residues that have shifted environments, reflect an evolutionary strategy to maintain receptor functionality while adapting to different biochemical landscapes. These changes, combined with the structural modification of removing transmembrane and intracellular domains, appear to have enabled the diversification of a soluble TLR5 form. This study provide insight into adaptation of specific environments (membranous and extracellular) and could guide additional computational analysis. Future studies could utilize molecular dynamics simulations for analyzing effects of environment-specific properties on protein conformational changes, as our previous studies shown intrinsic effects of structural shape on conformational characteristics on QTY-code designed water-soluble variants of membrane proteins (23). The findings of this study could also be utilized for designing algorithms to identify specific environmental adaptation pressures.

The strong evolutionary shift from phenylalanine (F) to tyrosine (Y) and arginine (R) to lysine (K) highlights the importance of polar and charged residues in molecular adaptation. By modulating the presence and distribution of such residues, protein engineers can design TLR5 variants or other immune receptors with tailored binding affinities, potentially improving the efficacy of immune responses or enhancing the specificity of receptor-ligand interactions which could have application in various therapeutic contexts. Understanding the evolutionary shifts from solvent to membrane environments and vice versa in TLR5 provides a framework for designing synthetic receptors that can adapt to different cellular environments. Our study suggests that utilizing variations with higher mutational barrier, especially acidic to basic (D>K) and polar to non-polar (T>V) changes, is potentially useful. It suggests a modular approach to protein evolution, where some regions largely coupled or sensitive to mutational changes across protein, while peripheral regions evolve more neutrally.

By mimicking evolutionary strategies that maintain functionality despite mutations, it may be possible to develop therapeutics that remain effective over time, even in the face of selective pressures such as drug resistance or immune evasion. The conserved nature of hydrophobic residues, alongside the strategic shifts in charged and polar residues, indicates the TLR5 receptor’s evolutionary adaptability to diverse biological roles, from membrane anchoring in its extracellular domain to soluble environments.

## Methods

### Protein sequence alignments and other characteristics

The protein sequence data were retrieved in FASTA format from Bai *et. al* (4). NCBI GenBank IDs was KM282522 and KR005612, respectively for mTLR5 and sTLR5. The membrane topology features of the proteins were also predicted using DeepTMHMM web application using the sequence data (13). The molecular weights (MW) and isoelectric points (pI) of the investigated proteins were calculated using the Expasy website (https://web.expasy.org/compute_pi/) (24, 25, 26).

Sequence alignment was generated using Clustsal Omaga (27) and visualized using Jalview 2.11.3.3 (28) and the alignments colour coded according to hydrophobicity. Each residue in the consensus sequence is determined as the most frequent residue in each column of the alignment. Sequence comparisons were done for the Clustal Omega aligned sequences (residues 1-626 for sTLR5 and 1-619 for mTLR5). Kyte-Doolittle scale was used as a measure of polarity od substituted residues (29). To compare the hydrophobicity of mTLR5 and sTLR5, we utilized the Kyte-Doolittle hydrophobicity scale for evaluating the aligned residues of both sequences. For each aligned residue, the hydrophobicity score was computed and plotted to visualize regional hydrophobicity differences between mTLR5 and sTLR5. A Python script (29, 30) was utilized to handle the data and generate plots utilizing BioPython 1.83 (31) and Matplotlib plotting library (32).

Codon usage bias is analyzed for the coding sequences (CDS), via GenRCA Rare Codon Analysis tool (https://www.genscript.com/tools/rare-codon-analysis) (33, 34, 35). Kazusa Codon Usage database for *Danio rerio* (zebrafish) were utilized as host organism. The codon adaptation index evaluates the relative efficiency of each codon by comparing them against a reference set of highly expressed genes within a species (33). The score for a gene is then determined based on the frequency with which these codons are used in that gene (33).

### Comparative structural analyses

Predictions for the structures of mTLR5 and sTLR5 were conducted through the AlphaFold3 program, accessed via (https://alphafoldserver.com/), following the instructions at the website (36). It employs deep learning techniques to predict protein structures with high accuracy (36). The structures were superposed using PyMOL Molecular Graphics System, version 3 (https://pymol.org/) (37). The similarities between structures analyzed quantitatively by the calculation of all-atom root <u>m</u>ean <u>s</u>quare <u>d</u>eviation (RMSD) values, no outlier reduction cycles were utilized (cycles=0).

### Evolutionary conservation profiles

The EVcouplings server (https://v2.evcouplings.org/) calculates <u>e</u>volutionary <u>c</u>ouplings (ECs) using a maximum entropy model (38), from the <u>m</u>ultiple <u>s</u>equence <u>a</u>lignments (MSAs). Traditional E-value thresholds can be inconsistent across different proteins due to variations in protein length and database size and the platform utilizes length-normalized bitscore thresholds instead, offering a more uniform measure of evolutionary distance (38, 39), For large sequence alignments, <u>P</u>seudo-likelihood <u>m</u>aximization (PLM) becomes asymptotically equivalent to full maximum-likelihood, resulting in more accurate contact predictions (39). In the context of this research, the EVcouplings-PLM method, which is the recommended approach, was applied to generate ECs for proteins.

The Consurf server (https://consurf.tau.ac.il/) was utilized to generate residue-wise evolutionary conservation profiles (40, 41, 42, 43), which was also used in our previous work (10, 11). The server processed aligned AlphaFold structures that were also used for RMSD calculations. Default settings were applied for homolog search, thresholds, alignment, phylogeny, and conservation scoring. Homologous sequences were retrieved from UniRef90, which clusters UniRef100 sequences at the 90 or 50% sequence identity levels using the MMseqs2 algorithm (44). ConSurf utilizes an MAFFT alignment algorithm to generate a multiple sequence alignment of the identified homologs. HMMER was employed for homolog identification with an E-value threshold of 0.0001. The search considered homologs with a maximal sequence identity of 95% and a minimal sequence identity of 35%. Conservation scores were determined using the Bayesian method, with the amino acid substitution model selected based on the best fit. The evolutionary conservation grades of each residue were visualized using PyMOL version 3 (37).

### Statistical Calculations

In this study, we analyzed the causal asymmetric relationships of evolutionary occurrence for individual amino acid in aligned residues. Our data is not in time continuous; hence it is hard to assume if the evolutionary dynamics was uniformly aggregate monotone (21). Since homoscedasticity assumptions may not met by our data, we used the generalCorr package in R (45, 46), which requires a kernel regression of x on y. Kernel regression is a non-parametric technique that can capture complex, non-linear relationships between variables without making strong assumptions about the underlying data distribution (45, 46). This package is particularly well-suited for our study because it does not rely on traditional parametric assumptions that our data might violate. The traditional Pearson correlation coefficients were also provided.

The observed correlations could be effect of a confounding variable that have nonlinear effect, the partial correlation coefficients (parcor_ijk) were calculated to assess the correlations after removing the nonlinear effects of the confounding variables (45, 46, 47). Evolutionary conservation is hypothesized as a possible confounding variable, since it may result in similarities of substitutions by reducing both frequencies. For the conservation estimations, ConSurf grades were used. ConSurf uses a phylogenetic approach, considering evolutionary history between sequences (40, 41, 42, 43). Causality is not symmetric, yet correlation coefficients are symmetric. The kernel causality was measured by comparing the absolute values of the two asymmetric correlations that were calculated using generalized (asymmetric) correlation coefficients (48). Statistical calculations were performed and visualized using R (The R Foundation for Statistical Computing, Vienna, Austria), version 4.4.0 (https://www.r-project.org/) (49).

## Ethics Approval

Ethics approval was not required for this computational study as it did not involve animal subjects, human participants, and identifiable data.

## Consent to participate

Not applicable. This computational study did not involve human participants.

## Consent for publication

Not applicable. This computational study did not involve human participants.

## Availability of data and materials

For more detailed information on the statistical analyses, input files and detailed outputs, including the AlphaFold3 calculations and codes to regenerate analyses, please visit the website: https://github.com/karagol-alper/TLR5soluble.

## Author Contributions

Authors contribute equally to this study. All authors have read and agreed to the published version of the manuscript.

## Acknowledgements

We thank Shuguang Zhang of the Massachusetts Institute of Technology (MIT) Media Lab for his valuable suggestions on presenting the data in Figure 1 and Tables.

## References

1) Kawai T, Ikegawa M, Ori D, et al. Decoding Toll-like receptors: Recent insights and perspectives in innate immunity. Immunity 2024;57(4):649–73.

2) Hayashi F, Smith KD, Ozinsky A, et al. The innate immune response to bacterial flagellin is mediated by Toll-like receptor 5. Nature 2001;410(6832):1099–103

3) Yoon S, Kurnasov O, Natarajan V, et al. Structural Basis of TLR5-Flagellin Recognition and Signaling. Science 2012;335(6070):859–64.

4) Bai JS, Li YW, Deng Y, et al. Molecular identification and expression analysis of TLR5M and TLR5S from orange-spotted grouper (Epinepheluscoioides). Fish & Shellfish Immunology 2017;63:97–102.

5) Jiang L, Pei L, Wang P, et al. Molecular Characterization and Evolution Analysis of Two Forms of TLR5 and TLR13 Genes Base on Larimichthys crocea Genome Data. International Journal of Genomics 2020;2020:1–17.

6) Wang Y, Zhang S, Li H, et al. Small-Molecule Modulators of Toll-like Receptors. Accounts of Chemical Research 2020;53(5):1046–55.

7) Jaenicke R. Protein stability and molecular adaptation to extreme conditons. European Journal of Biochemistry 1991;202(3):715–28.

8) Behzadi P, García-Perdomo HA, and Karpiński TM. Toll-like receptors: general molecular and structural biology. Journal of Immunology Research 2021;2021(1):9914854.

9) Ten Oever J, Kox M, van de Veerdonk FL, et al. The discriminative capacity of soluble Toll-like receptor (sTLR) 2 and sTLR4 in inflammatory diseases. BMC immunology 2014;15:1–0.

10) Karagöl A, Karagöl T, Smorodina E, et al. Structural bioinformatics studies of glutamate transporters and their AlphaFold2 predicted water-soluble QTY variants and uncovering the natural mutations of L-> Q, I-> T, F-> Y and Q-> L, T-> I and Y-> F. Plos one 2024;19(4):e0289644.

11) Karagöl T, Karagöl A, and Zhang S. Structural bioinformatics studies of serotonin, dopamine and norepinephrine transporters and their AlphaFold2 predicted water-soluble QTY variants and uncovering the natural mutations of L-> Q, I-> T, F-> Y and Q-> L, T-> I and Y-> F. Plos one 2024;19(3):e0300340.

12) Ruysschaert JM and Lonez C. Role of lipid microdomains in TLR-mediated signalling. Biochimica et Biophysica Acta (BBA)-Biomembranes 2015;1848(9):1860–7.

13) Hallgren J, Tsirigos KD, Pedersen MD, et al. DeepTMHMM predicts alpha and beta transmembrane proteins using deep neural networks. BioRxiv, 10 Apr 2022, preprint: not peer reviewed.

14) Gitlin I, Carbeck JD, and Whitesides GM. Why are proteins charged? Networks of charge– charge interactions in proteins measured by charge ladders and capillary electrophoresis. Angewandte Chemie International Edition 2006;45(19):3022–60.

15) Heads JT, Lamb R, Kelm S, et al. Electrostatic interactions modulate the differential aggregation propensities of IgG1 and IgG4P antibodies and inform charged residue substitutions for improved developability. *Protein Engineering*, Design and Selection 2019;32(6):277–88.

16) Zhang S, Tao F, Qing R, et al. QTY code enables design of detergent-free chemokine receptors that retain ligand-binding activities. Proceedings of the National Academy of Sciences 2018;115(37):E8652–9.

17) Wells L, Vosseller K, Cole RN, et al. Mapping sites of O-GlcNAc modification using affinity tags for serine and threonine post-translational modifications. Molecular & Cellular Proteomic 2002;1(10):791–804.

18) Wang ZA and Cole PA. The chemical biology of reversible lysine post-translational modifications. Cell chemical biology 2020;27(8):953–69.

19) Schwartz GW, Shauli T, Linial M, et al. Serine substitutions are linked to codon usage and differ for variable and conserved protein regions. Scientific reports 2019;9(1):17238.

20) Koonin EV. Splendor and misery of adaptation, or the importance of neutral null for understanding evolution. BMC biology 2016;14:1–8.

21) Cressman R. *Evolutionary dynamics and extensive form games*: MIT Press, 2003.

22) Gingold H and Pilpel Y. Determinants of translation efficiency and accuracy. Molecular systems biology 2011;7(1):481.

23) Karagöl A, Karagöl T, and Zhang S. Molecular dynamic simulations reveal that water-soluble QTY-Variants of glutamate transporters EAA1, EAA2 and EAA3 retain the conformational characteristics of native transporters. Pharmaceutical Research 2024;41(10):1965–77.

24) Gasteiger E, Hoogland C, Gattiker A, et al. Protein identification and analysis tools on the ExPASy server. Humana press 2005.

25) Bjellqvist B, Basse B, Olsen E, et al. Reference points for comparisons of two-dimensional maps of proteins from different human cell types defined in a pH scale where isoelectric points correlate with polypeptide compositions. Electrophoresis 1994;15(1):529–39.

26) Bjellqvist B, Hughes GJ, Pasquali C, et al. The focusing positions of polypeptides in immobilized pH gradients can be predicted from their amino acid sequences. Electrophoresis 1993;14(1):1023–31.

27) Sievers F and Higgins DG. Clustal Omega for making accurate alignments of many protein sequences. Protein Science 2018;27(1):135–45.

28) Waterhouse AM, Procter JB, Martin DM, et al. Jalview Version 2—a multiple sequence alignment editor and analysis workbench. Bioinformatics 2009;25(9):1189–91.

29) Kyte J and Doolittle RF. A simple method for displaying the hydropathic character of a protein. Journal of molecular biology 1982;157(1):105–32.

30) Karagöl A and Karagöl T. An Evolutionary Statistics Toolkit for Simplified Sequence Analysis on Web with Client-Side Processing. bioRxiv, 4 Aug 2024, preprint: not peer reviewed.

31) Cock PJ, Antao T, Chang JT, et al. Biopython: freely available Python tools for computational molecular biology and bioinformatics. Bioinformatics 2009;25(11):1422.

32) Hunter JD. Matplotlib: A 2D graphics environment. Computing in science & engineering 2007;9(03):90–5.

33) Sharp PM and Li WH. The codon adaptation index-a measure of directional synonymous codon usage bias, and its potential applications. Nucleic acids research 1987;15(3):1281–95.

34) Alexaki A, Kames J, Holcomb DD, et al. Codon and codon-pair usage tables (CoCoPUTs): facilitating genetic variation analyses and recombinant gene design. Journal of molecular biology 2019;431(13):2434–41.

35) Satapathy SS, Sahoo AK, Ray SK, et al. Codon degeneracy and amino acid abundance influence the measures of codon usage bias: improved Nc (N^c) and ENCprime (N^′ c) measures. Genes to Cells 2017;22(3):277–83.

36) Abramson J, Adler J, Dunger J, et al. Accurate structure prediction of biomolecular interactions with AlphaFold 3. Nature 2024:1–3.

37) Schrödinger LLC. The PyMOL Molecular Graphics System, Version 3, https://www.pymol.org. Accessed 17 September 2024

38) Hopf TA, Green AG, Schubert B, et al. The EVcouplings Python framework for coevolutionary sequence analysis. Bioinformatics 2019;35(9):1582–4.

39) Ekeberg M, Lövkvist C, Lan Y, et al. Improved contact prediction in proteins: using pseudolikelihoods to infer Potts models. *Physical Review E—Statistical*, Nonlinear, and Soft Matter Physics 2013;87(1):012707.

40) Yariv B, Yariv E, Kessel A, et al. Using evolutionary data to make sense of macromolecules with a “face-lifted” ConSurf. Protein Science 2023;32(3):e4582.

41) Ashkenazy H, Abadi S, Martz E, et al. ConSurf 2016: an improved methodology to estimate and visualize evolutionary conservation in macromolecules. Nucleic acids research 2016;44(W1):W344–50.

42) Celniker G, Nimrod G, Ashkenazy H, et al. ConSurf: using evolutionary data to raise testable hypotheses about protein function. Israel Journal of Chemistry 2013;53(3-4):199–206.

43) Landau M, Mayrose I, Rosenberg Y, et al. ConSurf 2005: the projection of evolutionary conservation scores of residues on protein structures. Nucleic acids research 2005;33(suppl_2):W299–302.

44) Suzek BE, Huang H, McGarvey P, et al. UniRef: comprehensive and non-redundant UniProt reference clusters. Bioinformatics 2007;23(10):1282–8.

45) Vinod HD. Generalized correlation and kernel causality with applications in development economics. Communications in Statistics-Simulation and Computation 2017;46(6):4513–34.

46) Vinod HD. Generalized correlations and kernel causality using R package generalCorr. SSRN 2016. Available at https://ssrn.com/abstract=2782223 or 10.2139/ssrn.2782223.

47) Kim S. ppcor: an R package for a fast calculation to semi-partial correlation coefficients. Communications for statistical applications and methods 2015;22(6):665.

48) Zheng S, Shi NZ, and Zhang Z. Generalized measures of correlation for asymmetry, nonlinearity, and beyond. Journal of the American Statistical Association 2012;107(499):1239–52.

49) R Core Team. R: A language and environment for statistical computing. Foundation for Statistical Computing, Vienna, Austria. 2013.

